# Effects of management on global crop pest damage depends on coevolutionary indicators

**DOI:** 10.64898/2026.04.10.717632

**Authors:** Hao Ran Lai, Jonathan D. Tonkin, Jason M. Tylianakis

**Affiliations:** Bioprotection Aotearoa Centre of Research Excellence, School of Biological Sciences, University of Canterbury, Christchurch 8140, Aotearoa New Zealand; South East Asia Rainforest Research Partnership, 88400 Kota Kinabalu, Sabah, Malaysia

## Abstract

Pests and pathogens reduce yields of major food crops by 15% globally^1^. Interventions such as pesticides can mitigate yield loss, but pests can evolve resistance to pesticides over short timescales^2,3^. Pest damage to crops may also be determined by (co)evolutionary selection on pests by crops themselves and vice versa^4^, yet it is unclear whether the relative evolutionary potential of pests versus crops predicts the outcome of this arms race. Here we test whether readily available indicators of the evolutionary potential^5–9^ of crops and their pests or pathogens influence global yield loss^10^ and its response to agricultural management^11–13^. We find that evolutionary potential of the pest (measured as its genome size) and crop (measured as population density or proximity of wild relatives) moderated the effect of agricultural management (seed importation, fertiliser and pesticide use) on crop damage. Crucially, statistical interactions among crop and pest evolutionary potentials explained as much variation in yield loss as did agricultural management. Our results show a huge spatial variability in management effectiveness and suggest greatest benefit in places with a stronger imbalance in the evolutionary potential between crop and pest. More broadly, our findings reveal a key role of evolution in determining present-day pest damage.

## Main

Globally, pests and pathogens (hereafter: “pests”) cause up to 15% of yield loss in five major food crops—wheat, rice, maize, potato and soybean—which constitute a quarter of the world’s agricultural land area, one-third of gross production value, and half of humanity’s daily caloric requirements^1,5^. Crop damage is expected to worsen with climate change^14^, reaching total crop loss where emergent pests strike unprepared production systems^15^. Given the growing human population is estimated to require a 56% increase in food production by 2050^16^, there is thus an urgent need to minimise yield loss.

Since the green revolution, pesticides and fertiliser have become indispensable for improving crop yield and combating pests^12,17^. However, chemical applications can often be excessive and suboptimal^18^, resulting in accelerated evolution of pesticide resistance^2,3^ alongside widespread pollution, eutrophication, non-target impacts^19–21^, climate impacts^22^, and benefits to some pests^23,24^. Crop breeding and genetic modification for pest resistance can also be overcome by pest counter-adaptations^4^. High-quality seed germplasm is imported to improve crop yield and pest resistance^25^, thereby reducing chemical reliance. Yet, poorly sanitised seed trade can worsen yield loss by spreading pathogens and facilitating evolutionary host switches^26,27^. Collectively, these conflicting patterns indicate substantial variability in the effectiveness of management actions for reducing yield loss, with outcomes often being driven by evolutionary mechanisms. In fact, agro-ecosystems are arenas for evolutionary arms races fought between the crop and pest, whereby their relative potential to evolve defensive traits or counterattack mechanisms, respectively, determines the net impact on crop yield loss^4,28,29^. Although the importance of evolution by natural selection (e.g., the need for pesticide regimes that avoid resistance) or artificial selection (e.g., plant breeding) are well recognised in food production, to our knowledge it remains unclear whether the evolutionary potentials of crops, pests and pathogens interact to explain global patterns of yield loss, or the extent to which these arms races are moderated by management practices.

Here we test the hypothesis that the influence of agricultural management practices on pest damage depends on the local evolutionary context, including crop and pest characteristics and their interaction with external selective forces^30,31^. Specifically, we test whether easily acquired indicators of the (co)evolutionary potential of crops, pests and pathogens influence yield loss, additively or interactively with conventional agricultural practices. As indicators, we hypothesise that pest species with large genomes would have a greater genetic repertoire with which to adapt to plant defenses^32–35^. For example, fungal pathogens with larger genomes are evolutionarily more versatile at overcoming crop defences or metabolising pesticides^32–35^, though virulent bacterial pathogens are often selected to have smaller genomes^36^. Although genome size in plants can arise from various mechanisms and may not correlate positively with evolutionary potential^37,38^, the genetic diversity of crops likely correlates positively with their population size^39,40^. and the availability of wild relatives as a source of new genetic material^41^. However, the alternative hypothesis is also plausible, as larger crop areas may also intensify selection on the pest, by offering fewer refuges to maintain susceptible pest populations^4^ and providing a greater reward for pests that overcome that crop’s defences. Similarly, adjacent wild relatives could provide pest-resistant genotypes to crops through sexual recombination^40,41^, or refuges to maintain vulnerable pest genotypes and prevent the fixation of pest virulence or pesticide/transgenic resistance in the population^42,43^. However, wild relatives could also act as a source of pests^44^, and selection on pests by different host plants can influence the evolution of pesticide resistance^45^. Moreover, native wild relatives can increase the number of native pests attacking crops^46^, thereby generating trade-offs by increasing the number of arms races in which the crop must engage. Consequently, competing pests can influence one another’s impact on the same crop, whereas a pest that specialises on a single crop faces stronger selection to overcome the crop’s defences^47,48^.

Importantly, the impact of a pest’s evolutionary potential likely depends on that of the crop, and the extent to which local conditions such as human management, allow the arms race to operate. For example, selection by pests on crops can only operate if seeds naturally disperse or are stored for use against the same pest populations in subsequent years. Conversely, repeated sowing of imported seed, which is not subject to selection by local pests, may ease pests’ evolutionary ability to overcome host-plant defences^15,49–51^. Similarly, fertilisation of crops can increase the reward for their consumption by pests, but also sustain stronger plant defences for the pest to overcome^52^. Meanwhile, although pesticides may select for pest resistance, this could be traded off against their ability to win biotic arms races against the plant^53,54^.

We frame these interactive effects as gene × gene × environment (𝐺 × 𝐺 × 𝐸) interactions, in-keeping with the coevolution literature^55^, though we make no assumptions about the specific gene(s) underpinning the arms race. To test our hypotheses, we use a global dataset^10^ of yield loss magnitude (hereafter: “yield loss”) for the world’s most important crops (wheat, rice, maize, potato and soybean) to 105 pest and pathogen taxa. Specifically, we test whether yield loss for a given crop–pest combination at a given location can be explained by available proxies of crop (𝐺_𝑐_, number of wild relative species per area; crop population density measured as harvest per area) and pest (𝐺_𝑝_, taxa-level genome size) coevolutionary potentials and their interactions with the management environment in which the (co)evolutionary arms races take place (𝐸fertiliser and pesticide use per area, seed import at national scale, and the biotic context of other competing arms races, which we measure as the relative number of arms races fought by the crop and pest; see *Methods*).

## Both management practices and evolutionary potentials determine yield loss

Crop yield loss to pests and pathogens was strongly determined by conventional agricultural management practices, as expected, but their impact depended equally on indicators of the evolutionary potentials of the crops, pests and pathogens themselves. The most parsimonious model retained all main and interaction terms (Table S1) and explained 37% of the variation in yield loss (44% when accounting for random effects). We further partitioned the explained variance by sets of predictors: the main effects of environment (which included well-known impacts of management) explained the highest proportion (11%); this was followed by the crop × pest × environment (𝐺 × 𝐺 × 𝐸) interactions (8%; Fig. 1b). The remaining variance in yield loss was explained by the two-way interactions (i.e., crop × pest, crop × environment, or pest × environment; 2–5% each), and even less was explained by the main effects of crop or pest evolutionary potential and background covariates (<2%). This ranking in variable importance was consistent whether all taxa were analysed together or separately (Fig. S1), suggesting that these impacts were not driven by systematic differences among pest taxa in genome size, which could have been confounded with other life-history differences among pest taxa.

**Figure 1.**
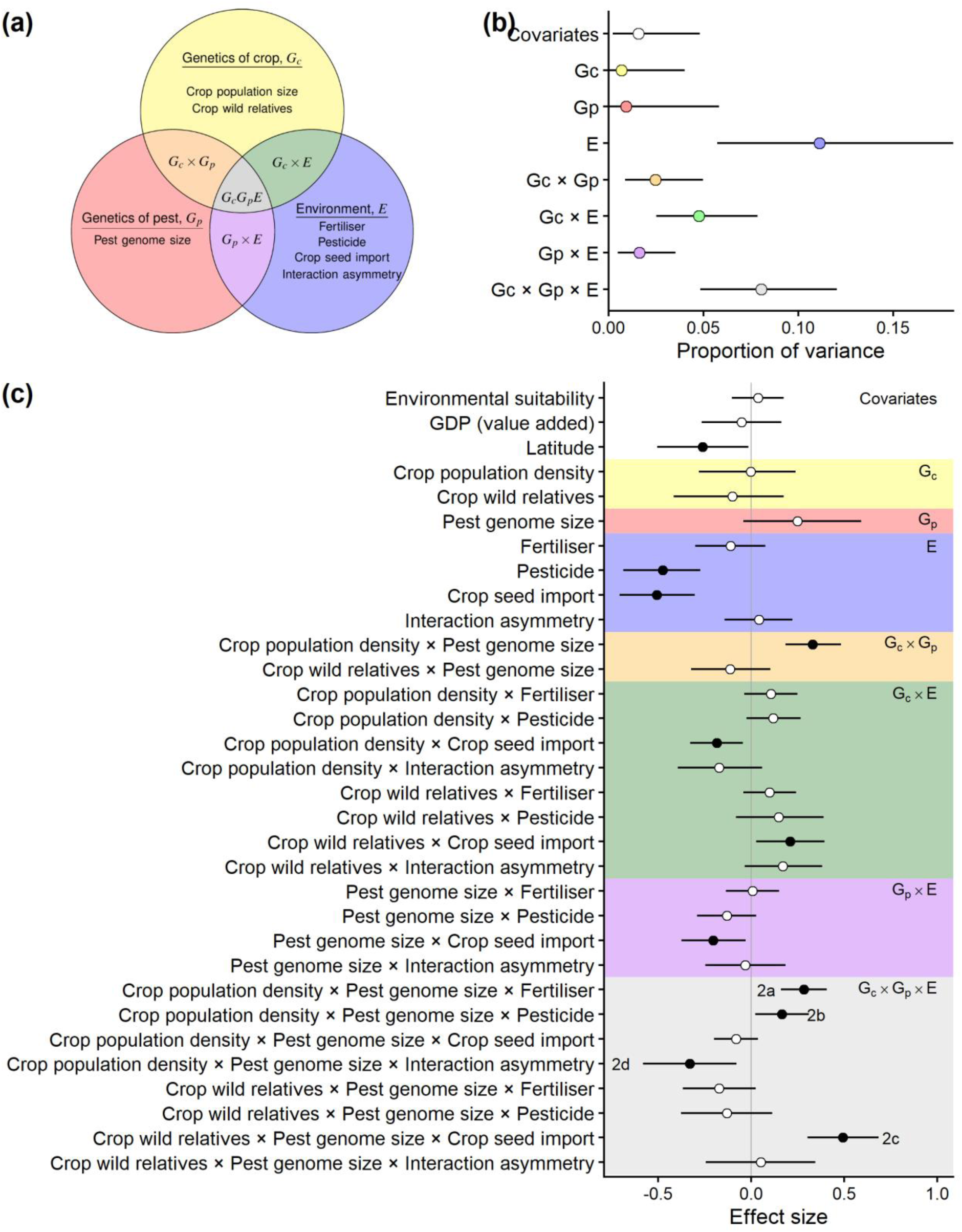
Variance components and effect sizes of the predictor sets. (a) Conceptual diagram showing the grouping of predictors and how they form interaction terms in our model. The same symbols and colours are used in panels b and c. The “covariates” group was not shown here, as it was not allowed to interact with other predictor sets. (b) Partitioning of total explained variance in yield loss magnitude by predictor groups. (c) Standardised effect size of model terms, quantifying their effect on yield loss magnitude (i.e. positive values increase yield loss to pests and pathogens). Points and error bars in b and c denote posterior median and 90% credible intervals. Open circles in panel c denote effects that overlap with zero. In the three-way gene × gene × environment interaction section (grey background), the strong interaction effects are annotated with the corresponding panels in Fig. 2a–d where each effect is plotted.

The main effects support existing evidence that yield loss declines with latitude^14^ and two agricultural management practices (i.e., the amount of pesticide^1^ and seed importation^51^). The lack of evidence for any main effect of crop or pest evolutionary potential on yield loss (main effects 𝐺_𝑐_ and 𝐺_𝑝_ in Fig. 1b,c) is expected, as the effect of pest evolutionary potential is likely to be relative to that of the crop, and vice versa. Indeed, crop and pest evolutionary potential had stronger effects on yield loss when they operated non-additively with one another (𝐺_𝑐_ × 𝐺_𝑝_ interactions), or when they were moderated by the environment (𝐺 × 𝐸 and 𝐺_𝑐_ × 𝐺_𝑝_ × 𝐸 interactions in Fig. 1c). With 17% of global yield-loss variation explained by indicators of crop and pest evolutionary potential and their interactions, crop management globally would benefit strongly from a more careful consideration of (co)evolution, either in the presence or absence of existing management practices.

Specifically, the 𝐺_𝑐_ × 𝐺_𝑝_ × 𝐸 interactions revealed how the impacts of management practices on yield loss are moderated or even reversed by crop and pest evolutionary potential (Fig. 2). Fertiliser, pesticide and seed import were more effective at reducing yield loss only when the arms race was asymmetric (i.e. pests and crops had contrasting evolutionary potentials, with one high and the other low; blue regions in Figs 2a–c), but these agricultural practices became less effective at reducing yield loss when crop and pest evolutionary potentials were comparable (i.e., both low or high; red regions in Figs 2a–c). This result indicates that, when an arms race is fought on more equal grounds between pests and crops, reducing agricultural inputs could be a strategy to alleviate economic and environmental costs without sacrificing crop yield. Our study therefore reveals potential coevolutionary mechanisms that may explain why farmers do not always need to spray more pesticide in larger crop fields^56–59^. Similarly, agricultural inputs may have greatest value in places with a stronger imbalance in the evolutionary potential between crop and pest. For example, when pests have a larger genome size and thus a broader genetic repertoire to rapidly adapt against the defences of host crops grown in small densities or around fewer wild relatives^32,34^ (top left corners of Figs 2a–c), applying fertiliser, pesticides or importing seeds seem to be more effective in reducing yield loss. However, this needs to be proceeded with caution because large-genome pests may also evolve pesticide resistance more rapidly^35^.

**Figure 2.**
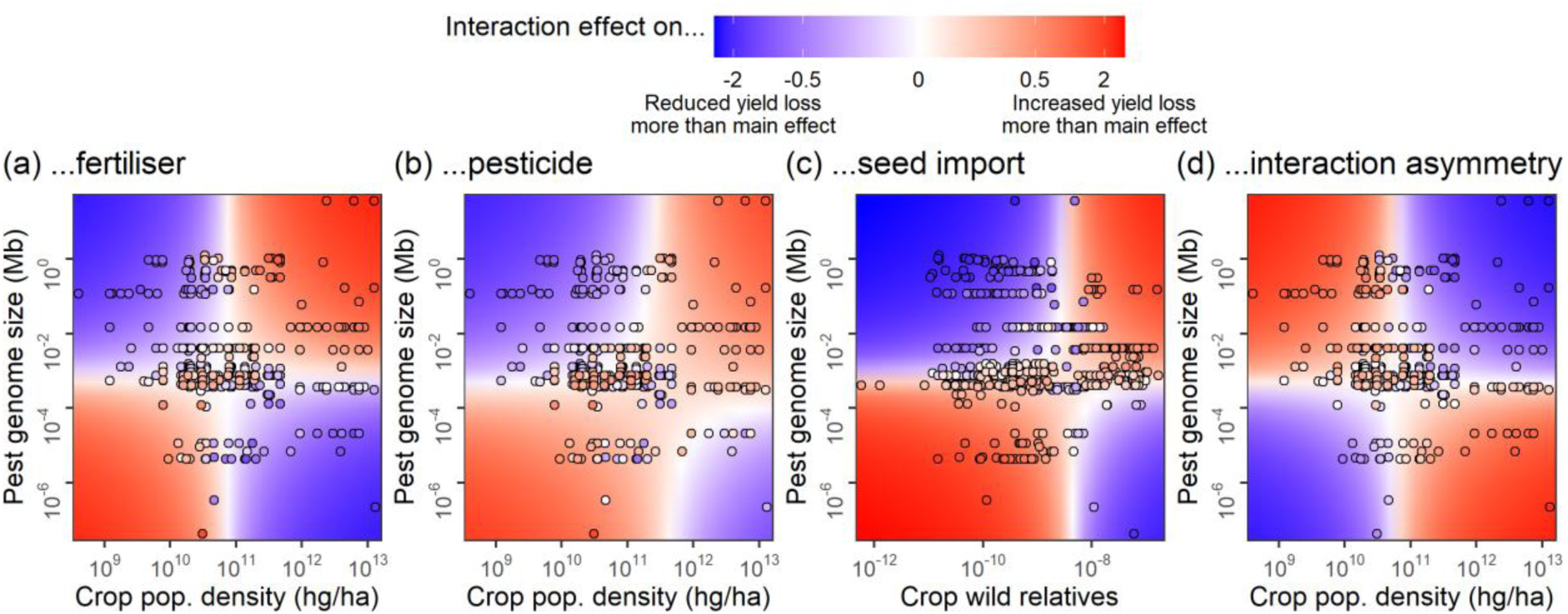
Gene × gene × environment interactions on yield loss magnitude. Each surface plot shows how the main effect of environment variables **(a)** fertiliser amount, **(b)** pesticide amount, **(c)** seed importation or **(d)** interaction asymmetry was moderated by indicator of pest and crop evolutionary potential (Y- and X-axis, respectively). Each colour surface was predicted by setting all other predictors at their means. Blue areas represent regions of evolutionary potential where the total effect reduced yield loss magnitude more than the main effect, whereas red areas represent the opposite. Filled circles are observed data, which are coloured by the fitted interaction effects (i.e., all predictors set to their observed values). All values are posterior medians.

## Suboptimal agricultural practices reveal huge potential for effectively allocating crop management resources

Although saving crop seed to replant locally could enable selection for resistance to pests^60^, we found that crops and regions with a greater seed importation suffered reduced yield loss (main effect of seed importation; Fig. 1c). Seed importation also eliminated the tendency for pests with larger genomes to cause more damage (seed importation × pest genome size interaction; Fig. 1c), potentially by disrupting a pest’s directional evolution. Unless imported seed is consistently sourced from a single location, temporal changes in crop genotypes would represent a moving target that interferes with a pest’s ability to adapt to each line’s chemical defences^25^, analogous to the fluctuating Red Queen dynamics that can make parasites less successful at attacking allopatric than sympatric host populations^61^. However, these benefits of seed importation diminished when there were many crop wild relatives in the landscape, suggesting that outcrossing with wild relatives may improve in situ crop evolution of pest and pathogen resistance^40,41^, thereby improving the success of local, relative to imported, seed^60^. Alternatively, the same pattern could arise from a different evolutionary mechanism, if the presence of wild relatives selects for host generalism in pests and pathogens^62^, thereby allowing them to exploit various imported crop lines.

When a generalist pest attacked a crop that was attacked by few other pest species (i.e., the pest was fighting more arms races than the crop), the pest’s effect on crop yield loss was greater when either pest genome size or crop population density was low and the other high (Fig. 2d). The lower resource availability of small crop areas likely selects for generalist pests that can maintain their populations on neighbouring crops or semi-natural habitats^63^, but that have the genetic repertoire to switch hosts^32–35^. This benefit of generalism is visible through the high yield loss on crops with small area caused by pests with large genome size when there is high interaction asymmetry. Pests with small genome size seem largely absent from crops with small populations (bottom-left corner of Fig. 2d), potentially because these pests have difficulty in fighting multiple arms races against different crops (as interaction asymmetry increases) and a crop with a small area may be insufficient to maintain populations of a specialist pest^63^.

Suboptimal impacts of management on yield loss against pests (i.e., when pesticide, fertiliser or seed importation did not reduce yield loss) were detected in about half of our data (49.7%), suggesting a huge potential for targeting crop management resources to where they can be more effective. Yet, resource allocation would need to be fine scaled, as globally there was huge spatial variation in the 𝐺_𝑐_ × 𝐺_𝑝_ × 𝐸 interactions, both within and among countries (Figs 3a–c). When these interaction effects were averaged by food-production regions and crop, none consistently benefited more or less from fertiliser, pesticide or seed import; only certain pest groups showed more consistent average interaction effects (Figs 3d–f). However, the average interaction effect obscures variation in 𝐺_𝑐_ × 𝐺_𝑝_ × 𝐸 dependencies within some groups. For example, hemipterans, maize and sub-Saharan Africa had a mean effect of 𝐺_𝑐_ × 𝐺_𝑝_ × 𝐸 interactions that was close to zero, but intervals that range from strongly positive to negative. This finding suggests that management decisions need to be specific to crop–pest combinations and made at the scale of a country or even smaller. Moreover, yield loss to pests is only one component of the economic trade-offs associated with agriculture, and yield gains from management such as fertiliser may offset the losses to pests.

**Figure 3.**
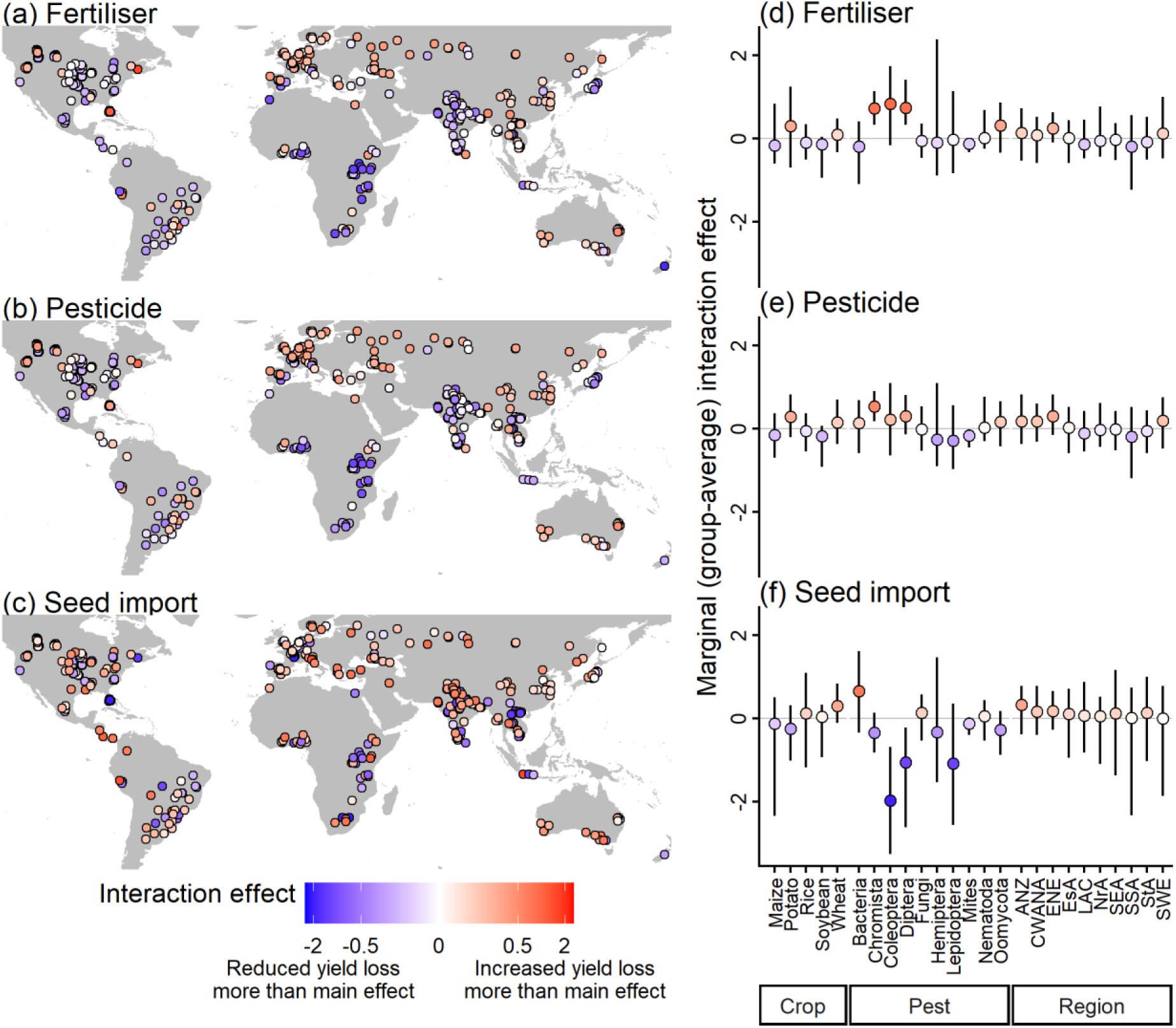
Geography (a-c) and marginal effects (d-f) of indicators of gene × gene × environment interaction effects on yield loss magnitude. Filled circles on maps are observed data, which are coloured by the posterior-median fitted interaction effects (i.e., all predictors set to their observed values). These fitted values are then broken down (marginalised across posteriors) by groups of crop, pest or region to show their group-average interactive effects on yield loss on the right (points and error bars are posterior median and 90% credible intervals, respectively). Key to regional abbreviations: ANZ = Australia and New Zealand; CWANA = Central-and-West Asia and North Africa; ENE = Eastern and Northern Europe; EsA = Eastern Asia; LAC = Latin America and Caribbean; NrA = Northern America; SEA = South-Eastern Asia; SSA = Sub-Saharan Africa; StA = Southern Asia; SWE = Southern and Western Europe.

Nevertheless, our results provide possible evolutionary explanations to findings of meta-analyses and case studies that showed huge variation in the effectiveness of pesticide or fertiliser across pests and crops^23,57,64^.

## Conclusion

Our findings offer evolutionary insights into where and why conventional agricultural practices reduce or exacerbate pest damage, which is crucial for the optimal distribution of management resources to improve food production while reducing their environmental costs^18,65^. Locations and taxa with the greatest variation, and therefore uncertainties, in management outcomes may merit further research and resource planning efforts^66^ (Fig. 3; see Table S2 for a lookup table). For example, increasing fertiliser addition alleviated the damage caused by the brown plant hopper on rice in Southern Asia, but exacerbated damage in Eastern Asia and varies from alleviating to exacerbating in South-Eastern Asia. Future research needs to uncover the underlying mechanisms that gave rise to the broad-scale emergent correlations detected in our study. Nevertheless, the explanatory power of evolutionary indicators and their interactions with management observed here suggests that relieving the burden of pathogens and pests on major food crops requires greater consideration of evolutionary biology in environmental decisions^67^. Crop management that overlooks these coevolutionary contexts may lead to wasteful actions or even exacerbate pest impacts^30,31,68^.

## Methods

### Datasets

We obtained measures of crop yield loss from a survey of crop health experts by Savary et al.^10^. In brief, the survey was hosted by the International Society for Plant Pathology (ISPP) through an online, worldwide questionnaire between 1 November 2016 and 31 January 2017. The questionnaire was designed to be simple to encourage responses^10^. For each of the five focal crops—wheat, rice, maize, potato and soybean—respondents were asked to answer four key questions: (1) approximate location where the crop and pest interacted, (2) pest or pathogen name (hereafter simply referred to as “pests”), (3) frequency of yield loss (recorded as chronic, frequent, infrequent, or rare), and (4) magnitude of yield loss due to the particular pest (recorded as <1%, 1–5%, 5–20%, 20–60%, or >60%). The survey resulted in 989 records from 219 experts in 67 countries after harmonisation^10^. We focused only on yield-loss magnitude in this study because yield-loss frequency was highly concentrated into one category and lacked variation; 70% of the survey responses were classified as “chronic”.

In this study, we sought to explain the magnitude of yield loss due to pests from the Savary et al.^10^ survey responses using sets of easily-measurable predictor variables that previous evidence suggests will likely correlate with the evolutionary potential of each crop and pest (i.e., which should relate to their additive genetic variance), and characteristics of the local environment (including management) that may influence crop-pest coevolution. The evolutionary variables were intended to capture each crop’s and pest’s intrinsic evolutionary potential, which may directly influence yield loss or influence yield loss non-additively by determining the outcome of arm races between crops and pests. Furthermore, yield loss is also known to be influenced by extrinsic selection drivers, which include agricultural practices or the impact of the surrounding crop/pest community on natural selection. In addition to the known main effects of these extrinsic factors, we are most interested in quantifying the strength of their interactions with crop and pest evolutionary potential.

Therefore, in the following analysis we selected proxies for three sets of predictors: crop evolutionary potential (𝐺_𝑐_), pest evolutionary potential (𝐺_𝑝_) and extrinsic factors (𝐸; hereafter simply referred to as “environment”); we then modelled yield loss as a function of their main effects, two-way interactions (analogous to gene × gene and gene × environment interactions), as well as three-way interactions (i.e., gene × gene × environment interactions; Fig. 1a). Note that the gene × gene × environment (𝐺 × 𝐺 × 𝐸) interaction terminology is used to reflect these concepts in the coevolution literature^55^, rather than implying the involvement of any specific gene(s).

For indicators of crop evolutionary potential (the predictor set 𝐺_𝑐_), we obtained harvested density (hg ha^−1^) by country from Food and Agriculture Organization of the United Nations^5^ as a proxy of population density^4,39,40^ and the richness of crop wild relative species (count ha ^−1^) as a proxy for the regional availability of genetic diversity or gene pool available to the crop through recombination^6,28^, though population density may alternatively indicate the potential resource available (reward) to pests that overcome its defences. We reproduced the distribution map of crop wild relatives by location following the methodology in Vincent et al.^6^ because the original published maps were no longer hosted on the authors’ website (see supplementary code).

For indicators of pest evolutionary potential (the predictor set 𝐺_𝑝_), we obtained each taxon’s genome size (Mb) from two databases. First, we standardised the pest taxonomic names using the R package taxize^69^ v0.9.100, and then extracted the genomic data from the Genomes on a Tree database^7^. For pests without any genome match, we further used the R package biomartr^8^ v1.0.5 to search from the National Center for Biotechnology Information (NCBI) database. We tried to retrieve genome size at the species level whenever possible and then filled in the remaining taxa with genus-level genome size. In the original yield loss survey data, some pests were reported as genera or species complex involving multiple taxa, for which we assigned genus-level or mean genome size across taxa.

For environmental selection (the predictor set 𝐸), we obtained proxies of both direct selection pressures due to management and moderators of natural selection, which we hypothesised could alter the outcome of crop-pest coevolution. For the key management drivers, we extracted the amount of nitrogen fertilizer input (kg ha^−1^) by location from Nishina et al.^11^. We then extracted crop-specific pesticide application (kg ha^−1^) by location from PEST-CHEMGRIDS^12^, and then matched the type of pesticide to the pest group (i.e., insecticide, fungicide, acaricide or nematicide). Thus, our analyses included pesticide application specific to the pest taxon for which yield loss was being predicted. In PEST-CHEMGRIDS^12^, potato was grouped with other vegetables into a larger group, so we approximated the potato-specific pesticide application by dividing the total pesticide application on vegetables by the local production area of potato obtained from Monfreda et al.^70^ (see supplementary code). Finally, we obtained the amount of seed import (USD) in 2016 by country from the World Integrated Trade Solution (WITS) database^71^, using the R package tradestatistics^72^ v 6.0.0 as a potential moderator of the potential for local crop–pest (co)evolution.

The surrounding community can also impose or modify biotic selective pressures^73^, and species that need to fight evolutionary arms races with many others may face trade-offs^47^. To capture this aspect of the biotic milieu, we used interaction asymmetry as a potential indicator of the relative intensity of selection pressure due to species interactions, which can influence coevolution^55^. To do so, we downloaded from the CABI Compendium^13^ a list of all plant species that were associated with every pest in our dataset, and then combined this with data on all the pests that attacked our five focal crops^10^ to construct a crop-by-pest adjacency matrix (i.e., interaction network). We used this matrix to calculate a metric for the relative number of interaction partners of each crop vs. each of its pests: interaction asymmetry^74^. A positive interaction asymmetry value indicates that the pest interacts with more plant species than the crop interacts with pest species (i.e., the pest has more interaction partners than the crop), whereas a negative value indicates the opposite. We hypothesise that negative asymmetry would advantage the pest over the crop, because selection on plant defensive traits will not occur unless those traits have correlated effects across the suite of pest species attacking a crop^75^. Instead, negative correlations among pests (particularly very different taxa such as pathogens and herbivores) may generate trade-offs for the plant^76^ and thereby slow the evolution of plant defences, in addition to potential non-additive ecological effects of multiple pests^47^. Additionally, while a generalist pest exerts strong selection pressure on a crop that interacts with fewer pests, it receives weaker selection reciprocally, such that generalist herbivores may be more vulnerable to plant defence compounds than specialists^77^. When interaction asymmetry is zero, it indicates that the pest and crop have equal numbers of interaction partners.

Finally, we included three background covariates that are known to be associated with yield loss. First, we extracted attainable yield (kg ha^−1^) during the period 2011–2040 from FAO’s GAEZ v4 database^78^ as a proxy of environmental suitability for each crop. Based on the assumption that more developed countries have more access to inputs to control yield loss^79^, we approximated the developmental status of each country’s agricultural sector using value added of the agriculture, forestry, and fishing industries (2016 US dollar) from the World Bank, using the R package wbstats^80^ v1.0.4. Lastly, we included the absolute value of latitude to control for the greater pest loadings on tropical than temperate crops^81^.

Prior to analysis, we excluded viruses (8% of yield loss data) from the pests, because they do not have pesticide data and they are often managed indirectly through measures to control their host vectors. We also removed two survey responses involving parasitic plants, because they do not form a sufficiently large group to model a random intercept. We further removed survey responses (22% of data) that did not have complete predictor data, resulting in 689 data points from 105 pest taxa and 53 countries. Prior to modelling, pest genome size, crop population density, number of crop wild relatives, fertiliser, pesticide, and value added of the agriculture industries were log-transformed, and we added 1 before log-transforming environmental suitability and crop seed import (to account for zero values). Then, all predictors were standardised to mean of zero and unit standard deviation to facilitate model convergence and the interpretation of slopes as partial effects.

### Statistical analyses

For each survey response 𝑖, we modelled the magnitude of yield loss, 𝑌_𝑖𝑐𝑝𝑟_, of crop 𝑐 due to pest 𝑝 in region 𝑟 as an ordinal response variable in a generalised linear mixed-effect model (GLMM) with cumulative distribution and logit link:

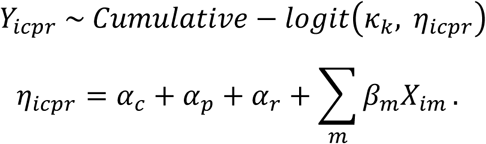

For an ordinal variable like yield loss magnitude, the cumulative-logit model^82^ estimates an underlying latent, continuous variable 𝜂 from which the 𝐾 = 5 ordinal scores were categorised and partitioned from 𝑘 = {1, 2, . . . , 𝐾 − 1} cutpoints, 𝜅_𝑘_.

In the linear predictor 𝜂, we began by including crop-, pest- and region-specific random intercepts as 𝛼_𝑐_, 𝛼_𝑝_ and 𝛼_𝑟_, respectively, to control for residual non-independence within taxa and regions. To estimate random intercepts, we treated each crop as a group (𝑐 = 1, 2, . . . , 5) in itself. For the pests, however, we classified them into ten larger taxonomic groups (𝑝 = 1, 2, . . . , 10) following Bebber et al.^81^, because over 42% of the pests were singletons in the dataset and estimating random intercepts based on one data point per pest would be problematic. Similarly, 21% of countries only had one data point, so we grouped the countries into ten food-production regions (𝑟 = 1, 2, . . . , 10) following the original study^10^ of yield loss, and included this region as a random intercept.

Next, we included predictors 𝑋 and their effects on yield loss magnitude 𝛽. In the full model, the predictors 𝑋 included the sets of main terms (𝐺_𝑐_, 𝐺_𝑝_ and 𝐸), two-way interaction terms (𝐺_𝑐_𝐺_𝑝_, 𝐺_𝑐_𝐸 and 𝐺_𝑝_𝐸), and three-way interaction terms (𝐺_𝑐_𝐺_𝑝_𝐸), as listed in Fig. 1a. For parsimony, we constructed a series of candidate models (see Table S1) that correspond to a few broad classes of hypothesis: yield loss was only associated with the background covariates (Model 1); yield loss was also additively influenced by genetics and the environment (Models 2–7); yield loss was also non-additively influenced by indicators of evolutionary potential and environment, but restricted to two-way interactions (Model 8–17); and finally the full model that also contained three-way interactions (Model 18). We compared the candidate models using the Leave-One-Out Information Criterion (LOOIC) from the loo package^83^ v2.5.1 and then selected the best-supported model with the lowest LOOIC value. In addition to the main analysis above, we also performed a set of sensitivity analyses to examine if the effects were different when analysed separately by crop or by pest. We subset the full data by five crop (maize, potato, rice, soybean, and wheat) or three pest groups (arthropods, bacteria, and fungi and oomycetes) and fitted the same 18 candidate models.

Each candidate model was fitted via Bayesian inference using the brms package^84^ v2.20.1 in R^85^ v4.2.1, with 1,000 Hamiltonian Monte Carlo (HMC) warmups and 1,000 post-warmup samplings across four chains, resulting in 4,000 posterior samples in total. To promote model convergence, we tuned the adaptive HMC step size to 0.99. Chain convergence was assessed with the Gelman–Rubin diagnostic (𝑅^ < 1.05). We diagnosed the model residuals using a quantile–quantile plot (Fig. S2).

To compare variable importance, we used part 𝑅^2^ to partition the proportion of variance in yield loss explained by each predictor set 𝐺_𝑐_, 𝐺_𝑝_ and 𝐸, as well as their interaction terms^86^. To examine the variation of 𝐸 (i.e., fertiliser, pesticide and seed importation) across crops, pests and regions, we also calculated their realised effects as the combination of main and interaction effects. In simpler terms, the total effect of 𝐸 is 𝛽_𝐸_𝐸 + 𝛽_𝐸𝐺𝑐_𝐸𝐺_𝑐_ + 𝛽_𝐸𝐺𝑝_𝐸𝐺_𝑝_ + 𝛽_𝐸𝐺𝑐𝐺𝑝_𝐸𝐺_𝑐_𝐺_𝑝_, which can be rearranged to emphasise how crop and pest evolutionary potentials moderate the main effect of agricultural inputs: (𝛽_𝐸_ + [𝛽_𝐸𝐺𝑐_𝐺_𝑐_ + 𝛽_𝐸𝐺𝑝_𝐺_𝑝_ + 𝛽_𝐸𝐺𝑐𝐺𝑝_𝐺_𝑐_𝐺_𝑝_]) 𝐸. We used the terms in the square brackets to quantify how much of the main effect of an agricultural input was increased (exacerbated crop yield loss) or decreased (alleviated crop yield loss) due to crop and pest evolutionary potentials.

## Acknowledgements and funding statement

This research, and HRL, JDT and JMT were supported by the Bioprotection Aotearoa Centre of Research Excellence. HRL was supported by the Marsden Fund Council from New Zealand Government, funding managed by the Royal Society Te Apārangi (grant MFP-UOC2102). JDT was supported by a Rutherford Discovery Fellowship administered by the Royal Society Te Apārangi (RDF-18-UOC-007). JMT was funded by the Ministry for Business, Innovation and Employment (programmes C09X2104 and C09X2209).

## Author Contributions

HRL, JDT and JMT conceived the research. HRL compiled data, performed analyses and drafted the manuscript. All authors interpreted the results and wrote the final article.

