## Supplementary Information for "Effects of management on global crop pest damage depends on coevolutionary indicators"

#### Comparison of candidate models

Table S1: Candidate models using all data containing different sets of predictors (see Fig. 1 in the main text) and the difference in LOOIC from the best model, which is Model 18 in bold. Key to predictor sets: $C$ = background covariates including latitude, environmental suitability and gross domestic production value; $G_c$ = crop evolutionary potential including population density and wild relative species richness; $G_p$ = pest evolutionary potential including genome size; $E$ = environmental factors including fertiliser, pesticide, crop seed import and crop-pest interaction asymmetry.

| Model | Predictor set | | | | | | | | ΔLOOIC |
|---|---|---|---|---|---|---|---|---|---|
| | $C$ | $G_c$ | $G_p$ | $E$ | $G_cG_p$ | $G_cE$ | $G_pE$ | $G_cG_pE$ | |
| 1 | ✓ | | | | | | | | 27.0 |
| 2 | ✓ | ✓ | | | | | | | 24.4 |
| 3 | ✓ | | ✓ | | | | | | 28.6 |
| 4 | ✓ | | | ✓ | | | | | 16.2 |
| 5 | ✓ | ✓ | ✓ | | | | | | 26.6 |
| 6 | ✓ | ✓ | | ✓ | | | | | 17.0 |
| 7 | ✓ | | ✓ | ✓ | | | | | 16.6 |
| 8 | ✓ | ✓ | ✓ | | ✓ | | | | 22.4 |
| 9 | ✓ | ✓ | | ✓ | | ✓ | | | 13.4 |
| 10 | ✓ | | ✓ | ✓ | | | ✓ | | 20.0 |
| 11 | ✓ | ✓ | ✓ | ✓ | | | | | 18.4 |
| 12 | ✓ | ✓ | ✓ | ✓ | ✓ | | | | 16.2 |
| 13 | ✓ | ✓ | ✓ | ✓ | | ✓ | | | 14.4 |
| 14 | ✓ | ✓ | ✓ | ✓ | | | ✓ | | 23.0 |
| 15 | ✓ | ✓ | ✓ | ✓ | ✓ | ✓ | | | 23.0 |
| 16 | ✓ | ✓ | ✓ | ✓ | ✓ | | ✓ | | 21.2 |
| 17 | ✓ | ✓ | ✓ | ✓ | | ✓ | ✓ | | 18.6 |
| **18** | ✓ | ✓ | ✓ | ✓ | ✓ | ✓ | ✓ | ✓ | 0.0 |

### Crop-, pest- and region-specific effects of agricultural inputs

Table S2: Realised effect of agricultural inputs on the damage of pest and pathogen species on each of the five major crops by region. The crops, pest and pathogen, and region represents all of the analysed data from the survey by the International Society for Plant Pathology (ISPP; Savary et al. 2019). The realised effect of agricultural inputs are the effects accounting for the context dependencies on the crop's and pest's evolutionary potentials, including crop population density, crop wild relative species richness, and pest genome size. Positive and negative realised effects indicate increased and decreased yield loss more than expected, respectively. Values are reported as posterior median with 90% credible intervals in parentheses.

| | | | Effect of agricultural inputs | | |
|---|---|---|---|---|---|
| Crop | Pest or pathogen | Region | Fertiliser | Pesticide | Seed import |
| Maize | African boll worm | Sub Saharan Africa | -0.04 (-0.55, 0.46) | -0.46 (-1.06, 0.14) | -1.70 (-2.41, -1.06) |
| | African stem borer | Sub Saharan Africa | -0.54 (-1.13, 0.32) | -0.74 (-1.33, -0.09) | -1.14 (-2.78, -0.28) |
| | Anthracnose leaf blight | Northern America | -0.19 (-0.49, 0.12) | -0.25 (-0.77, 0.21) | -0.20 (-0.52, 0.11) |
| | Anthracnose stalk rot | Northern America | -0.13 (-0.40, 0.14) | -0.16 (-0.59, 0.23) | -0.22 (-0.48, 0.03) |
| | Asian stem borer | South-Eastern Asia | 0.04 (-0.27, 0.33) | -0.32 (-0.67, 0.03) | -1.17 (-1.62, -0.73) |
| | Bacterial Stalk Rot | Southern Asia | 0.10 (-0.32, 0.53) | 0.17 (-0.34, 0.70) | 1.25 (0.69, 1.81) |
| | Banded Leaf and Sheath Blight | Southern Asia | -0.14 (-0.33, 0.04) | -0.23 (-0.48, 0.00) | 0.13 (-0.09, 0.34) |
| | Black Maize Beetle | Sub Saharan Africa | 0.04 (-0.68, 0.75) | -0.51 (-1.34, 0.30) | -2.51 (-3.49, -1.62) |
| | Brown stripe downy mildew | Southern Asia | -0.23 (-0.46, -0.02) | -0.41 (-0.68, -0.14) | -0.46 (-0.78, -0.15) |
| | Common rust | Latin America and Caribbean | -0.16 (-0.40, 0.08) | -0.34 (-0.70, -0.01) | -0.39 (-0.71, -0.08) |
| | | Northern America | 0.06 (-0.18, 0.31) | -0.10 (-0.46, 0.22) | -0.59 (-0.92, -0.33) |
| | | Southern Asia | -0.19 (-0.39, -0.01) | -0.33 (-0.57, -0.09) | -0.15 (-0.42, 0.09) |
| | Common smut | Latin America and Caribbean | -0.23 (-0.42, -0.04) | -0.05 (-0.32, 0.21) | 0.38 (0.17, 0.59) |
| | | Northern America | -0.45 (-0.82, -0.09) | -0.25 (-0.82, 0.30) | 0.55 (0.19, 0.92) |
| | Corn stunt | Latin America and Caribbean | -0.24 (-0.73, 0.26) | 0.15 (-0.46, 0.76) | 1.79 (1.10, 2.47) |
| | Crazy top | Latin America and Caribbean | -0.06 (-0.27, 0.14) | -0.28 (-0.55, -0.01) | -0.62 (-0.92, -0.34) |
| | Cutworm | Sub Saharan Africa | 0.01 (-0.63, 0.65) | -0.49 (-1.23, 0.25) | -2.21 (-3.08, -1.40) |
| | Diabrotica | Northern America | 0.92 (0.12, 1.72) | 0.07 (-0.85, 0.95) | -2.78 (-3.93, -1.83) |
| | | Southern & Western Europe | 0.82 (0.18, 1.47) | 0.07 (-0.63, 0.77) | -2.35 (-3.17, -1.60) |
| | | Southern Asia | 0.14 (-0.40, 0.66) | -0.38 (-0.99, 0.23) | -2.00 (-2.76, -1.31) |
| | Diplodia rot | Latin America and Caribbean | -0.13 (-0.30, 0.02) | -0.14 (-0.39, 0.11) | 0.27 (0.07, 0.46) |
| | | Northern America | -0.26 (-0.55, 0.02) | -0.16 (-0.63, 0.25) | 0.11 (-0.16, 0.37) |
| | Ear rot | Southern Asia | -0.12 (-0.27, 0.02) | -0.11 (-0.34, 0.10) | 0.25 (0.07, 0.42) |
| | European stem borer | Northern America | 0.76 (0.17, 1.36) | 0.08 (-0.57, 0.73) | -2.14 (-2.89, -1.45) |
| | | Southern & Western Europe | 0.68 (0.06, 1.31) | 0.00 (-0.70, 0.70) | -2.21 (-3.08, -1.47) |
| | Eyespot | Northern America | -0.37 (-0.71, -0.04) | -0.25 (-0.80, 0.26) | 0.40 (0.05, 0.72) |
| | | Southern & Western Europe | -0.41 (-0.79, -0.03) | -0.31 (-0.92, 0.27) | 0.45 (0.06, 0.82) |
| | F&G ear rots | Eastern Asia | -0.12 (-0.28, 0.02) | -0.11 (-0.39, 0.10) | 0.07 (-0.09, 0.23) |
| | | Latin America and Caribbean | -0.16 (-0.42, 0.03) | -0.21 (-0.65, 0.07) | 0.16 (-0.09, 0.43) |
| | | Northern America | -0.19 (-0.48, 0.08) | -0.16 (-0.64, 0.23) | -0.06 (-0.32, 0.20) |
| | | Southern & Western Europe | -0.18 (-0.44, 0.07) | -0.15 (-0.59, 0.21) | -0.04 (-0.28, 0.19) |
| | | Southern Asia | -0.17 (-0.37, 0.03) | -0.22 (-0.56, 0.09) | 0.14 (-0.11, 0.37) |
| | | Sub Saharan Africa | -0.10 (-0.33, 0.13) | -0.20 (-0.46, 0.08) | 0.33 (0.06, 0.59) |
| | F&G stalk rots | Latin America and Caribbean | -0.14 (-0.39, 0.03) | -0.18 (-0.60, 0.07) | 0.16 (-0.07, 0.41) |
| | | Northern America | -0.20 (-0.49, 0.08) | -0.18 (-0.66, 0.23) | -0.06 (-0.33, 0.21) |
| | | Southern & Western Europe | -0.17 (-0.44, 0.07) | -0.15 (-0.59, 0.22) | -0.05 (-0.29, 0.19) |
| | | Southern Asia | -0.16 (-0.36, 0.05) | -0.23 (-0.54, 0.07) | 0.19 (-0.07, 0.46) |
| | | Sub Saharan Africa | -0.10 (-0.33, 0.13) | -0.19 (-0.45, 0.08) | 0.33 (0.07, 0.59) |
| | Fall armyworm | Northern America | 0.83 (0.33, 1.34) | 0.14 (-0.40, 0.67) | -1.84 (-2.47, -1.27) |
| | | Sub Saharan Africa | -0.68 (-1.10, -0.28) | -0.81 (-1.30, -0.33) | -0.83 (-1.45, -0.23) |
| | Head smut | Southern & Western Europe | -0.35 (-0.65, -0.04) | -0.20 (-0.68, 0.25) | 0.36 (0.05, 0.65) |
| | Maize stem borer | Sub Saharan Africa | 0.01 (-0.63, 0.64) | -0.49 (-1.22, 0.25) | -2.16 (-3.02, -1.37) |
| | Mediterranean corn borer | Southern & Western Europe | 0.88 (0.23, 1.53) | 0.11 (-0.59, 0.82) | -2.34 (-3.15, -1.59) |
| | Northern leaf blight | Eastern Asia | -0.21 (-0.45, 0.03) | -0.25 (-0.67, 0.13) | 0.08 (-0.20, 0.35) |
| | | Latin America and Caribbean | -0.10 (-0.35, 0.03) | -0.10 (-0.54, 0.09) | 0.10 (-0.09, 0.37) |
| | | Northern America | -0.18 (-0.45, 0.08) | -0.15 (-0.59, 0.23) | -0.07 (-0.31, 0.17) |

Table S2: Realised effect of agricultural inputs on the damage of pest and pathogen species on each of the five major crops by region. The crops, pest and pathogen, and region represents all of the analysed data from the survey by the International Society for Plant Pathology (ISPP; Savary et al. 2019). The realised effect of agricultural inputs are the effects accounting for the context dependencies on the crop's and pest's evolutionary potentials, including crop population density, crop wild relative species richness, and pest genome size. Positive and negative realised effects indicate increased and decreased yield loss more than expected, respectively. Values are reported as posterior median with 90% credible intervals in parentheses. *(continued)*

| | | | Effect of agricultural inputs | | |
|---|---|---|---|---|---|
| Crop | Pest or pathogen | Region | Fertiliser | Pesticide | Seed import |
| | | Southern & Western Europe | -0.18 (-0.42, 0.06) | -0.16 (-0.56, 0.20) | -0.03 (-0.27, 0.19) |
| | | Southern Asia | -0.12 (-0.33, 0.07) | -0.20 (-0.48, 0.06) | 0.25 (0.01, 0.49) |
| | | Sub Saharan Africa | -0.11 (-0.35, 0.13) | -0.22 (-0.50, 0.07) | 0.33 (0.05, 0.60) |
| | Root knot nematodes | Sub Saharan Africa | -0.24 (-0.53, 0.05) | -0.35 (-0.83, 0.09) | 0.14 (-0.21, 0.48) |
| | Silk fly | Northern America | 0.86 (0.32, 1.42) | 0.14 (-0.45, 0.73) | -2.00 (-2.68, -1.37) |
| | Sorghum downy mildew | Southern Asia | -0.23 (-0.46, -0.02) | -0.41 (-0.68, -0.14) | -0.46 (-0.78, -0.15) |
| | Southern leaf blight | Eastern Asia | -0.25 (-0.52, 0.02) | -0.26 (-0.71, 0.17) | 0.27 (-0.03, 0.55) |
| | | Latin America and Caribbean | -0.22 (-0.49, 0.04) | -0.28 (-0.72, 0.13) | 0.36 (0.03, 0.68) |
| | | Southern Asia | -0.12 (-0.34, 0.10) | -0.19 (-0.49, 0.11) | 0.38 (0.11, 0.65) |
| | Southern rust | Eastern Asia | -0.06 (-0.31, 0.17) | -0.25 (-0.61, 0.08) | -0.51 (-0.81, -0.21) |
| | | Latin America and Caribbean | -0.09 (-0.23, 0.04) | -0.21 (-0.41, -0.02) | -0.22 (-0.42, -0.04) |
| | | Northern America | 0.08 (-0.19, 0.33) | -0.08 (-0.48, 0.24) | -0.60 (-0.93, -0.34) |
| | | Southern Asia | -0.19 (-0.39, -0.01) | -0.33 (-0.57, -0.09) | -0.15 (-0.42, 0.09) |
| | | Sub Saharan Africa | -0.25 (-0.46, -0.04) | -0.36 (-0.61, -0.13) | -0.00 (-0.28, 0.26) |
| | Tar Spot | Latin America and Caribbean | -0.21 (-0.46, 0.04) | -0.28 (-0.69, 0.10) | 0.27 (-0.04, 0.57) |
| | White spot | Latin America and Caribbean | -0.29 (-0.79, 0.15) | -0.04 (-0.72, 0.55) | 1.47 (0.87, 2.20) |
| Potato | Aphids | Southern & Western Europe | 1.08 (0.58, 1.56) | 0.65 (0.23, 1.07) | -0.62 (-1.14, -0.09) |
| | Brown rot | Central and West Asia and North Africa | -0.93 (-1.60, -0.24) | -0.05 (-0.80, 0.70) | 0.14 (-0.41, 0.68) |
| | | Sub Saharan Africa | -0.20 (-0.69, 0.22) | 0.25 (-0.20, 0.71) | 0.14 (-0.42, 0.54) |
| | Canker and black scurf | Latin America and Caribbean | 0.17 (-0.15, 0.49) | 0.35 (-0.01, 0.68) | -0.31 (-0.65, 0.03) |
| | Colorado potato beetle | Northern America | 1.45 (0.84, 2.07) | 0.75 (0.18, 1.32) | -1.33 (-2.00, -0.72) |
| | | Southern & Western Europe | 1.15 (0.64, 1.64) | 0.59 (0.15, 1.03) | -0.89 (-1.41, -0.39) |
| | Common scab | Central and West Asia and North Africa | -0.49 (-1.06, -0.01) | 0.09 (-0.47, 0.59) | 0.09 (-0.30, 0.51) |
| | | Latin America and Caribbean | -0.56 (-1.19, 0.07) | 0.17 (-0.54, 0.86) | -0.28 (-0.82, 0.23) |
| | | Northern America | -0.75 (-1.43, -0.07) | 0.06 (-0.70, 0.80) | -0.16 (-0.71, 0.38) |
| | | Southern & Western Europe | -0.76 (-1.44, -0.07) | 0.05 (-0.70, 0.79) | -0.14 (-0.69, 0.40) |
| | | Southern Asia | -0.57 (-1.03, -0.08) | 0.06 (-0.47, 0.58) | 0.11 (-0.29, 0.49) |
| | | Sub Saharan Africa | -0.96 (-1.60, -0.29) | -0.18 (-0.97, 0.57) | 0.39 (-0.15, 0.92) |
| | Cyst nematode | Central and West Asia and North Africa | 0.25 (0.03, 0.50) | 0.24 (0.03, 0.47) | -0.32 (-0.64, -0.06) |
| | | Southern & Western Europe | 0.39 (0.05, 0.75) | 0.41 (0.08, 0.74) | -0.46 (-0.84, -0.07) |
| | Early blight | Central and West Asia and North Africa | -0.05 (-0.30, 0.20) | 0.20 (-0.08, 0.48) | -0.12 (-0.36, 0.12) |
| | | Latin America and Caribbean | -0.06 (-0.44, 0.33) | 0.29 (-0.15, 0.72) | -0.28 (-0.65, 0.08) |
| | | Northern America | -0.10 (-0.53, 0.33) | 0.27 (-0.22, 0.73) | -0.32 (-0.72, 0.08) |
| | | Southern & Western Europe | -0.10 (-0.50, 0.28) | 0.23 (-0.20, 0.65) | -0.25 (-0.62, 0.10) |
| | | Southern Asia | -0.12 (-0.43, 0.18) | 0.15 (-0.20, 0.47) | -0.13 (-0.39, 0.13) |
| | | Sub Saharan Africa | -0.13 (-0.54, 0.28) | 0.22 (-0.25, 0.65) | -0.27 (-0.63, 0.10) |
| | Late blight | Central and West Asia and North Africa | 0.41 (0.16, 0.65) | 0.28 (0.07, 0.49) | -0.30 (-0.57, -0.00) |
| | | Eastern & Northern Europe | 0.45 (0.18, 0.93) | 0.32 (0.08, 0.76) | -0.30 (-0.64, 0.02) |
| | | Eastern Asia | 0.43 (0.16, 0.71) | 0.34 (0.08, 0.59) | -0.18 (-0.48, 0.14) |
| | | Latin America and Caribbean | 0.57 (0.24, 0.94) | 0.39 (0.09, 0.73) | -0.51 (-0.88, -0.09) |
| | | Northern America | 0.71 (0.32, 1.10) | 0.52 (0.19, 0.86) | -0.57 (-0.99, -0.14) |
| | | Southern & Western Europe | 0.65 (0.28, 1.05) | 0.48 (0.16, 0.83) | -0.47 (-0.88, -0.05) |
| | | Southern Asia | 0.48 (0.20, 0.76) | 0.28 (0.02, 0.56) | -0.60 (-0.98, -0.10) |
| | | Sub Saharan Africa | 0.15 (-0.08, 0.80) | 0.11 (-0.13, 0.51) | -0.03 (-1.04, 0.31) |
| | Leaf miner | Latin America and Caribbean | 0.59 (0.27, 0.91) | 0.32 (0.04, 0.59) | -0.38 (-0.71, -0.05) |
| | Leaf worm | Southern Asia | 1.25 (0.71, 1.78) | 0.52 (0.02, 1.00) | -1.15 (-1.73, -0.62) |
| | Potato leafhopper | Northern America | 2.74 (1.54, 3.95) | 1.30 (0.15, 2.42) | -1.07 (-2.19, 0.03) |
| | | Southern & Western Europe | 1.84 (0.90, 2.78) | 0.75 (-0.13, 1.60) | -0.61 (-1.47, 0.24) |
| | | Sub Saharan Africa | 2.61 (1.53, 3.70) | 1.08 (0.00, 2.18) | -2.43 (-3.59, -1.40) |

Table S2: Realised effect of agricultural inputs on the damage of pest and pathogen species on each of the five major crops by region. The crops, pest and pathogen, and region represents all of the analysed data from the survey by the International Society for Plant Pathology (ISPP; Savary et al. 2019). The realised effect of agricultural inputs are the effects accounting for the context dependencies on the crop's and pest's evolutionary potentials, including crop population density, crop wild relative species richness, and pest genome size. Positive and negative realised effects indicate increased and decreased yield loss more than expected, respectively. Values are reported as posterior median with 90% credible intervals in parentheses. *(continued)*

| | | | Effect of agricultural inputs | | |
|---|---|---|---|---|---|
| Crop | Pest or pathogen | Region | Fertiliser | Pesticide | Seed import |
| | Powdery scab | Australia and New Zealand | 0.73 (0.32, 1.15) | 0.59 (0.23, 0.97) | -0.37 (-0.82, 0.13) |
| | | Northern America | 0.72 (0.32, 1.11) | 0.53 (0.20, 0.88) | -0.51 (-0.94, -0.08) |
| | Verticillium wilt | Central and West Asia and North Africa | -0.08 (-0.34, 0.17) | 0.18 (-0.11, 0.46) | -0.10 (-0.33, 0.14) |
| | | Northern America | -0.21 (-0.69, 0.26) | 0.19 (-0.35, 0.69) | -0.27 (-0.67, 0.13) |
| | | Southern & Western Europe | -0.11 (-0.45, 0.24) | 0.22 (-0.17, 0.60) | -0.19 (-0.51, 0.12) |
| | | Sub Saharan Africa | -0.17 (-0.60, 0.25) | 0.20 (-0.28, 0.65) | -0.25 (-0.62, 0.12) |
| | Zebra chip | Australia and New Zealand | -1.86 (-3.16, -0.53) | -0.26 (-1.71, 1.16) | -0.34 (-1.40, 0.66) |
| Rice | Aggregate sheath spot | Northern America | -0.14 (-0.39, 0.10) | -0.22 (-0.63, 0.16) | -0.18 (-0.45, 0.07) |
| | Bacterial blight | Eastern Asia | -0.48 (-1.43, -0.04) | -0.03 (-1.25, 0.50) | 1.23 (0.52, 2.77) |
| | | Northern America | -0.49 (-0.83, -0.14) | -0.01 (-0.44, 0.40) | 0.94 (0.53, 1.36) |
| | | South-Eastern Asia | -0.10 (-0.57, 0.47) | 0.19 (-0.33, 0.68) | 0.82 (0.20, 1.59) |
| | | Southern Asia | 0.00 (-0.34, 0.33) | 0.20 (-0.20, 0.58) | 0.91 (0.40, 1.50) |
| | | Sub Saharan Africa | 0.35 (-0.07, 0.82) | 0.39 (-0.08, 0.92) | 0.69 (0.18, 1.20) |
| | Bacterial panicle blight | Latin America and Caribbean | -0.10 (-0.72, 0.39) | 0.14 (-0.41, 0.51) | 0.72 (0.29, 1.42) |
| | | Southern Asia | 0.10 (-0.17, 0.38) | 0.27 (-0.03, 0.59) | 0.49 (0.14, 0.85) |
| | Bacterial sheath rot | South-Eastern Asia | 0.19 (-0.14, 0.51) | 0.33 (-0.03, 0.72) | 0.44 (0.05, 0.83) |
| | Bakanae | Eastern Asia | -0.07 (-0.18, 0.03) | -0.04 (-0.20, 0.11) | 0.03 (-0.08, 0.13) |
| | Blast | Eastern Asia | -0.17 (-0.54, 0.01) | -0.11 (-0.70, 0.16) | 0.19 (-0.01, 0.53) |
| | | Latin America and Caribbean | -0.14 (-0.39, 0.02) | -0.08 (-0.48, 0.13) | 0.20 (0.02, 0.44) |
| | | Northern America | -0.19 (-0.49, 0.01) | -0.11 (-0.60, 0.18) | 0.17 (-0.04, 0.45) |
| | | South-Eastern Asia | -0.04 (-0.23, 0.12) | 0.02 (-0.26, 0.18) | 0.20 (0.05, 0.40) |
| | | Southern & Western Europe | -0.28 (-0.73, 0.01) | -0.27 (-0.98, 0.20) | 0.27 (-0.06, 0.70) |
| | | Southern Asia | -0.07 (-0.46, 0.06) | -0.05 (-0.62, 0.12) | 0.27 (0.09, 0.61) |
| | | Sub Saharan Africa | 0.01 (-0.20, 0.20) | 0.03 (-0.25, 0.26) | 0.26 (0.08, 0.46) |
| | Brown plant hopper | Eastern Asia | 0.31 (0.02, 0.60) | -0.12 (-0.42, 0.18) | -1.10 (-1.50, -0.71) |
| | | South-Eastern Asia | 0.05 (-0.53, 0.38) | -0.27 (-0.69, 0.05) | -0.64 (-1.47, 0.02) |
| | | Southern Asia | -0.15 (-0.43, 0.11) | -0.39 (-0.70, -0.08) | -0.64 (-1.15, 0.10) |
| | Brown spot | Latin America and Caribbean | -0.10 (-0.21, 0.01) | -0.04 (-0.21, 0.12) | 0.22 (0.07, 0.35) |
| | | Northern America | -0.14 (-0.27, 0.01) | -0.02 (-0.22, 0.17) | 0.14 (-0.01, 0.27) |
| | | South-Eastern Asia | -0.02 (-0.25, 0.12) | 0.04 (-0.29, 0.18) | 0.23 (0.06, 0.46) |
| | | Southern & Western Europe | -0.19 (-0.51, 0.01) | -0.16 (-0.61, 0.16) | 0.29 (0.04, 0.53) |
| | | Southern Asia | -0.05 (-0.18, 0.07) | -0.03 (-0.18, 0.12) | 0.27 (0.12, 0.43) |
| | False smut | Eastern Asia | -0.05 (-0.15, 0.04) | -0.04 (-0.20, 0.10) | -0.03 (-0.13, 0.06) |
| | | Southern Asia | -0.06 (-0.15, 0.02) | -0.08 (-0.20, 0.03) | 0.11 (-0.00, 0.21) |
| | Kernel smut | Southern Asia | -0.03 (-0.19, 0.13) | 0.05 (-0.13, 0.23) | 0.43 (0.21, 0.65) |
| | Leaf folder | South-Eastern Asia | 0.16 (-0.11, 0.43) | -0.22 (-0.52, 0.09) | -1.09 (-1.49, -0.69) |
| | Narrow brown leaf spot | Northern America | -0.10 (-0.23, 0.03) | -0.02 (-0.21, 0.16) | 0.06 (-0.07, 0.18) |
| | Rice sheath mite | Southern Asia | -0.07 (-0.15, 0.00) | -0.13 (-0.23, -0.04) | -0.05 (-0.15, 0.05) |
| | Sheath blight | Eastern Asia | -0.04 (-0.14, 0.05) | -0.04 (-0.19, 0.09) | -0.06 (-0.16, 0.03) |
| | | Latin America and Caribbean | -0.06 (-0.14, 0.02) | -0.07 (-0.21, 0.05) | -0.01 (-0.11, 0.08) |
| | | Northern America | -0.06 (-0.26, 0.09) | -0.07 (-0.41, 0.14) | -0.12 (-0.31, 0.02) |
| | | South-Eastern Asia | -0.01 (-0.11, 0.04) | -0.01 (-0.12, 0.07) | 0.09 (0.01, 0.23) |
| | | Southern Asia | -0.04 (-0.13, 0.03) | -0.05 (-0.17, 0.05) | 0.10 (0.00, 0.18) |
| | Sheath rot | Eastern Asia | -0.09 (-0.20, 0.02) | -0.03 (-0.20, 0.12) | 0.09 (-0.03, 0.20) |
| | | Northern America | -0.10 (-0.22, 0.03) | -0.02 (-0.20, 0.16) | 0.05 (-0.07, 0.17) |
| | | South-Eastern Asia | 0.01 (-0.09, 0.12) | 0.07 (-0.07, 0.18) | 0.15 (0.03, 0.31) |
| | | Southern & Western Europe | -0.16 (-0.35, 0.02) | -0.18 (-0.50, 0.10) | 0.20 (-0.03, 0.41) |
| | | Southern Asia | -0.05 (-0.15, 0.05) | -0.04 (-0.16, 0.08) | 0.20 (0.07, 0.33) |
| | Stem borers | Eastern Asia | 0.32 (0.02, 0.61) | -0.12 (-0.43, 0.19) | -1.15 (-1.55, -0.74) |

Table S2: Realised effect of agricultural inputs on the damage of pest and pathogen species on each of the five major crops by region. The crops, pest and pathogen, and region represents all of the analysed data from the survey by the International Society for Plant Pathology (ISPP; Savary et al. 2019). The realised effect of agricultural inputs are the effects accounting for the context dependencies on the crop's and pest's evolutionary potentials, including crop population density, crop wild relative species richness, and pest genome size. Positive and negative realised effects indicate increased and decreased yield loss more than expected, respectively. Values are reported as posterior median with 90% credible intervals in parentheses. *(continued)*

| | | | Effect of agricultural inputs | | |
|---|---|---|---|---|---|
| Crop | Pest or pathogen | Region | Fertiliser | Pesticide | Seed import |
| | | Northern America | 0.44 (0.13, 0.76) | -0.04 (-0.36, 0.28) | -1.19 (-1.60, -0.78) |
| | | South-Eastern Asia | 0.02 (-0.52, 0.39) | -0.27 (-0.70, 0.08) | -0.43 (-1.49, 0.12) |
| | | Southern & Western Europe | 0.42 (-0.33, 1.07) | -0.20 (-1.00, 0.52) | -2.24 (-3.11, -1.48) |
| | | Southern Asia | -0.13 (-0.40, 0.13) | -0.40 (-0.72, -0.09) | -0.80 (-1.34, -0.35) |
| | | Sub Saharan Africa | -0.42 (-0.76, -0.07) | -0.51 (-0.92, -0.12) | -0.11 (-0.53, 0.31) |
| | Stem rot | Northern America | -0.28 (-0.57, 0.01) | -0.24 (-0.72, 0.20) | 0.23 (-0.06, 0.52) |
| | White Grubs | South-Eastern Asia | -0.01 (-0.34, 0.30) | -0.38 (-0.77, -0.01) | -1.24 (-1.74, -0.75) |
| Soybean | Anthracnose | Latin America and Caribbean | -0.11 (-0.26, 0.03) | -0.17 (-0.37, 0.00) | 0.12 (-0.06, 0.28) |
| | Armyworm | Northern America | -0.38 (-0.70, -0.06) | -0.54 (-0.93, -0.17) | -0.43 (-0.88, 0.01) |
| | | Southern & Western Europe | -0.33 (-0.66, -0.02) | -0.57 (-0.96, -0.19) | -0.94 (-1.44, -0.43) |
| | Brown Spot | Latin America and Caribbean | -0.06 (-0.20, 0.08) | -0.09 (-0.25, 0.09) | 0.26 (0.09, 0.43) |
| | | Northern America | -0.02 (-0.12, 0.08) | -0.00 (-0.11, 0.11) | 0.21 (0.09, 0.33) |
| | Cercospora leaf blight | Northern America | -0.01 (-0.14, 0.12) | 0.02 (-0.12, 0.17) | 0.29 (0.12, 0.45) |
| | Charcoal rot | Latin America and Caribbean | -0.08 (-0.21, 0.05) | -0.12 (-0.27, 0.04) | 0.17 (0.02, 0.33) |
| | Cylindrocladium rot | Eastern Asia | -0.01 (-0.26, 0.25) | -0.09 (-0.36, 0.20) | 0.39 (0.13, 0.66) |
| | Cyst nematode | Latin America and Caribbean | -0.13 (-0.28, 0.02) | -0.20 (-0.40, -0.01) | 0.08 (-0.11, 0.26) |
| | | Northern America | -0.06 (-0.16, 0.02) | -0.08 (-0.23, 0.01) | 0.09 (-0.03, 0.18) |
| | Downy mildew | Latin America and Caribbean | -0.18 (-0.32, -0.03) | -0.30 (-0.46, -0.13) | -0.25 (-0.47, -0.05) |
| | Frogeye leaf spot | Northern America | -0.02 (-0.13, 0.08) | -0.01 (-0.14, 0.11) | 0.21 (0.08, 0.34) |
| | | Sub Saharan Africa | 0.00 (-0.36, 0.36) | -0.14 (-0.52, 0.27) | 0.50 (0.15, 0.86) |
| | Fusarium wilt and rot | Latin America and Caribbean | -0.09 (-0.21, 0.04) | -0.13 (-0.28, 0.02) | 0.14 (-0.00, 0.29) |
| | Phomopsis seed decay | Latin America and Caribbean | -0.05 (-0.20, 0.10) | -0.07 (-0.25, 0.11) | 0.30 (0.11, 0.48) |
| | Phytophthora root & stem rot | Eastern Asia | -0.40 (-0.62, -0.16) | -0.46 (-0.69, -0.22) | -0.03 (-0.33, 0.27) |
| | | Latin America and Caribbean | -0.19 (-0.35, -0.03) | -0.30 (-0.53, -0.12) | -0.25 (-0.61, 0.01) |
| | | Northern America | -0.14 (-0.27, -0.01) | -0.24 (-0.40, -0.10) | -0.20 (-0.48, 0.02) |
| | | Sub Saharan Africa | -0.60 (-0.93, -0.27) | -0.61 (-0.93, -0.29) | 0.15 (-0.25, 0.55) |
| | Pythium damping-off | Latin America and Caribbean | -0.19 (-0.38, -0.02) | -0.35 (-0.58, -0.12) | -0.40 (-0.68, -0.14) |
| | | Northern America | -0.13 (-0.26, -0.01) | -0.23 (-0.38, -0.09) | -0.19 (-0.42, 0.03) |
| | Reniform nematode | Northern America | -0.02 (-0.11, 0.07) | -0.01 (-0.10, 0.09) | 0.17 (0.07, 0.27) |
| | Rhizoctonia rot and blight | Latin America and Caribbean | -0.07 (-0.24, 0.06) | -0.10 (-0.35, 0.05) | 0.15 (-0.04, 0.29) |
| | | Northern America | -0.07 (-0.17, 0.03) | -0.10 (-0.23, 0.02) | 0.11 (-0.02, 0.23) |
| | | Southern Asia | -0.18 (-0.47, 0.11) | -0.31 (-0.62, 0.02) | 0.32 (-0.00, 0.63) |
| | | Sub Saharan Africa | -0.12 (-0.41, 0.17) | -0.20 (-0.49, 0.11) | 0.40 (0.12, 0.68) |
| | Root knot nematodes | Northern America | -0.01 (-0.11, 0.10) | 0.01 (-0.09, 0.13) | 0.21 (0.09, 0.33) |
| | | Sub Saharan Africa | 0.06 (-0.21, 0.33) | 0.00 (-0.29, 0.31) | 0.37 (0.12, 0.62) |
| | Soybean rust | Eastern Asia | -0.59 (-0.91, -0.27) | -0.65 (-1.01, -0.31) | -0.33 (-0.78, 0.10) |
| | | Latin America and Caribbean | -0.26 (-0.52, -0.01) | -0.45 (-0.75, -0.15) | -0.64 (-1.17, -0.09) |
| | | Northern America | -0.18 (-0.39, 0.05) | -0.34 (-0.61, -0.10) | -0.40 (-1.09, -0.06) |
| | | Southern Asia | -0.75 (-1.22, -0.29) | -0.87 (-1.40, -0.36) | -0.81 (-1.70, -0.10) |
| | | Sub Saharan Africa | -0.99 (-1.60, -0.50) | -0.99 (-1.61, -0.47) | -0.43 (-1.21, 0.29) |
| | Spider mites | Latin America and Caribbean | -0.16 (-0.33, 0.00) | -0.28 (-0.50, -0.06) | -0.12 (-0.34, 0.10) |
| | Stem canker | Latin America and Caribbean | -0.05 (-0.20, 0.10) | -0.07 (-0.25, 0.11) | 0.30 (0.11, 0.48) |
| | Sudden death | Latin America and Caribbean | -0.08 (-0.21, 0.05) | -0.12 (-0.27, 0.04) | 0.18 (0.02, 0.33) |
| | | Northern America | -0.07 (-0.20, 0.04) | -0.08 (-0.28, 0.05) | 0.15 (0.01, 0.28) |
| | Target spot | Latin America and Caribbean | -0.12 (-0.30, 0.06) | -0.19 (-0.44, 0.06) | 0.25 (0.02, 0.47) |
| | | Northern America | -0.05 (-0.15, 0.06) | -0.05 (-0.16, 0.07) | 0.19 (0.07, 0.31) |
| | White mold | Latin America and Caribbean | -0.11 (-0.27, 0.05) | -0.17 (-0.38, 0.04) | 0.18 (-0.01, 0.38) |
| | | Northern America | -0.06 (-0.18, 0.05) | -0.08 (-0.24, 0.06) | 0.16 (0.02, 0.29) |
| Wheat | Aphids | Eastern Asia | -0.05 (-0.63, 0.54) | -0.03 (-0.67, 0.61) | 0.92 (0.32, 1.52) |

Table S2: Realised effect of agricultural inputs on the damage of pest and pathogen species on each of the five major crops by region. The crops, pest and pathogen, and region represents all of the analysed data from the survey by the International Society for Plant Pathology (ISPP; Savary et al. 2019). The realised effect of agricultural inputs are the effects accounting for the context dependencies on the crop's and pest's evolutionary potentials, including crop population density, crop wild relative species richness, and pest genome size. Positive and negative realised effects indicate increased and decreased yield loss more than expected, respectively. Values are reported as posterior median with 90% credible intervals in parentheses. *(continued)*

| | | | Effect of agricultural inputs | | |
|---|---|---|---|---|---|
| Crop | Pest or pathogen | Region | Fertiliser | Pesticide | Seed import |
| | | Northern America | -0.27 (-0.73, 0.20) | -0.24 (-0.78, 0.27) | 0.64 (0.15, 1.11) |
| | | Southern & Western Europe | -0.10 (-0.91, 0.55) | -0.06 (-0.82, 0.60) | 0.93 (0.26, 1.91) |
| | | Southern Asia | -0.37 (-0.83, 0.09) | -0.32 (-0.86, 0.19) | 0.63 (0.14, 1.12) |
| | | Sub Saharan Africa | -0.84 (-1.42, -0.24) | -0.63 (-1.32, 0.03) | 0.99 (0.35, 1.64) |
| | Army worm | Southern Asia | -0.42 (-0.93, 0.11) | -0.38 (-0.99, 0.19) | 0.68 (0.13, 1.20) |
| | Aster Yellows | Northern America | 0.95 (-0.32, 2.22) | 1.18 (-0.41, 2.77) | -1.13 (-2.35, 0.07) |
| | Crown and Root Rot | Australia and New Zealand | 0.11 (-0.15, 0.37) | 0.17 (-0.21, 0.56) | 0.31 (0.07, 0.55) |
| | | Eastern Asia | 0.15 (-0.05, 0.36) | 0.29 (-0.03, 0.64) | 0.09 (-0.15, 0.33) |
| | | Northern America | 0.18 (-0.09, 0.46) | 0.32 (-0.11, 0.78) | 0.19 (-0.10, 0.48) |
| | False armyworm | Sub Saharan Africa | -0.28 (-0.80, 0.25) | -0.28 (-0.88, 0.30) | 0.68 (0.13, 1.21) |
| | FHB-scab | Central and West Asia and North Africa | 0.14 (-0.10, 0.37) | 0.21 (-0.14, 0.58) | 0.21 (-0.01, 0.44) |
| | | Eastern & Northern Europe | 0.15 (-0.08, 0.38) | 0.28 (-0.09, 0.65) | 0.12 (-0.16, 0.40) |
| | | Eastern Asia | 0.15 (-0.06, 0.38) | 0.28 (-0.06, 0.66) | 0.09 (-0.16, 0.35) |
| | | Latin America and Caribbean | 0.09 (-0.08, 0.27) | 0.16 (-0.10, 0.43) | 0.18 (0.02, 0.35) |
| | | Northern America | 0.12 (-0.07, 0.34) | 0.22 (-0.08, 0.56) | 0.15 (-0.05, 0.36) |
| | | Southern & Western Europe | 0.16 (-0.07, 0.41) | 0.32 (-0.04, 0.71) | 0.04 (-0.23, 0.34) |
| | Heterodera avenae | Northern America | 0.08 (-0.14, 0.31) | 0.20 (-0.13, 0.57) | 0.32 (0.08, 0.59) |
| | Heterodera filipjevi | Northern America | 0.08 (-0.14, 0.31) | 0.20 (-0.13, 0.57) | 0.32 (0.08, 0.59) |
| | Leaf rust | Australia and New Zealand | -0.09 (-0.37, 0.20) | 0.03 (-0.34, 0.42) | 0.57 (0.28, 0.87) |
| | | Central and West Asia and North Africa | -0.07 (-0.37, 0.22) | 0.03 (-0.34, 0.43) | 0.50 (0.22, 0.84) |
| | | Eastern & Northern Europe | 0.09 (-0.11, 0.30) | 0.18 (-0.09, 0.48) | 0.25 (0.01, 0.50) |
| | | Eastern Asia | 0.09 (-0.09, 0.26) | 0.16 (-0.08, 0.42) | 0.22 (0.01, 0.44) |
| | | Northern America | -0.00 (-0.18, 0.19) | 0.07 (-0.17, 0.36) | 0.31 (0.11, 0.56) |
| | | Southern & Western Europe | 0.09 (-0.17, 0.33) | 0.20 (-0.14, 0.54) | 0.32 (0.01, 0.69) |
| | | Southern Asia | -0.04 (-0.22, 0.14) | 0.02 (-0.21, 0.27) | 0.32 (0.14, 0.51) |
| | Nodorum blotch | Australia and New Zealand | 0.24 (-0.10, 0.59) | 0.31 (-0.20, 0.86) | 0.21 (-0.11, 0.52) |
| | | Central and West Asia and North Africa | 0.20 (-0.08, 0.50) | 0.29 (-0.13, 0.76) | 0.16 (-0.11, 0.43) |
| | | Eastern & Northern Europe | 0.17 (-0.08, 0.42) | 0.30 (-0.08, 0.69) | 0.08 (-0.22, 0.36) |
| | | Eastern Asia | 0.19 (-0.08, 0.46) | 0.27 (-0.13, 0.69) | 0.17 (-0.09, 0.42) |
| | | Latin America and Caribbean | 0.11 (-0.07, 0.30) | 0.18 (-0.09, 0.46) | 0.17 (-0.01, 0.34) |
| | | Northern America | 0.16 (-0.06, 0.40) | 0.27 (-0.08, 0.65) | 0.12 (-0.12, 0.34) |
| | | Southern & Western Europe | 0.19 (-0.06, 0.48) | 0.32 (-0.08, 0.78) | 0.07 (-0.19, 0.35) |
| | Powdery mildew | Eastern Asia | 0.13 (-0.08, 0.35) | 0.26 (-0.06, 0.60) | 0.23 (-0.02, 0.51) |
| | | Southern & Western Europe | 0.17 (-0.03, 0.37) | 0.29 (-0.00, 0.60) | 0.12 (-0.13, 0.40) |
| | Pratylenchus neglectus | Northern America | 0.36 (-0.08, 0.81) | 0.51 (-0.14, 1.16) | -0.18 (-0.59, 0.24) |
| | Pratylenchus thornei | Australia and New Zealand | 0.41 (-0.02, 0.85) | 0.44 (-0.17, 1.05) | 0.04 (-0.37, 0.44) |
| | | Northern America | 0.34 (-0.07, 0.77) | 0.49 (-0.13, 1.11) | -0.15 (-0.55, 0.26) |
| | Rhizoctonia Bare patch | Australia and New Zealand | 0.08 (-0.19, 0.36) | 0.16 (-0.23, 0.59) | 0.36 (0.11, 0.63) |
| | Russian wheat aphid | Sub Saharan Africa | -0.21 (-0.64, 0.22) | -0.19 (-0.69, 0.28) | 0.59 (0.13, 1.03) |
| | Sharp Eye Spot | Eastern Asia | 0.12 (-0.05, 0.28) | 0.23 (-0.03, 0.51) | 0.11 (-0.08, 0.32) |
| | Spot blotch | Central and West Asia and North Africa | 0.21 (-0.07, 0.49) | 0.29 (-0.12, 0.72) | 0.14 (-0.13, 0.40) |
| | | Eastern & Northern Europe | 0.19 (-0.08, 0.46) | 0.32 (-0.09, 0.73) | 0.05 (-0.26, 0.35) |
| | | Northern America | 0.22 (-0.06, 0.51) | 0.35 (-0.10, 0.81) | 0.06 (-0.22, 0.35) |
| | | Southern & Western Europe | 0.22 (-0.07, 0.51) | 0.37 (-0.08, 0.83) | 0.02 (-0.27, 0.32) |
| | | Southern Asia | 0.18 (-0.06, 0.43) | 0.26 (-0.10, 0.64) | 0.13 (-0.11, 0.37) |
| | Stem rust | Australia and New Zealand | -0.09 (-0.37, 0.19) | 0.02 (-0.34, 0.40) | 0.56 (0.28, 0.85) |
| | | Central and West Asia and North Africa | -0.05 (-0.28, 0.19) | 0.05 (-0.26, 0.39) | 0.45 (0.20, 0.71) |
| | | Eastern & Northern Europe | 0.01 (-0.22, 0.21) | 0.09 (-0.20, 0.37) | 0.33 (0.09, 0.62) |
| | | Eastern Asia | -0.04 (-0.27, 0.20) | 0.06 (-0.25, 0.39) | 0.46 (0.21, 0.71) |
| | | Northern America | -0.00 (-0.18, 0.19) | 0.06 (-0.17, 0.35) | 0.30 (0.11, 0.55) |

Table S2: Realised effect of agricultural inputs on the damage of pest and pathogen species on each of the five major crops by region. The crops, pest and pathogen, and region represents all of the analysed data from the survey by the International Society for Plant Pathology (ISPP; Savary et al. 2019). The realised effect of agricultural inputs are the effects accounting for the context dependencies on the crop's and pest's evolutionary potentials, including crop population density, crop wild relative species richness, and pest genome size. Positive and negative realised effects indicate increased and decreased yield loss more than expected, respectively. Values are reported as posterior median with 90% credible intervals in parentheses. *(continued)*

| | | | Effect of agricultural inputs | | |
|---|---|---|---|---|---|
| Crop | Pest or pathogen | Region | Fertiliser | Pesticide | Seed import |
| | | Southern & Western Europe | 0.00 (-0.28, 0.29) | 0.14 (-0.25, 0.56) | 0.54 (0.23, 0.86) |
| | | Southern Asia | -0.02 (-0.23, 0.19) | 0.07 (-0.22, 0.37) | 0.39 (0.17, 0.62) |
| | | Sub Saharan Africa | -0.05 (-0.26, 0.16) | 0.03 (-0.25, 0.32) | 0.40 (0.19, 0.62) |
| | Stripe rust | Australia and New Zealand | -0.10 (-0.36, 0.17) | -0.00 (-0.34, 0.36) | 0.53 (0.27, 0.80) |
| | | Central and West Asia and North Africa | -0.04 (-0.30, 0.25) | 0.08 (-0.27, 0.49) | 0.52 (0.23, 0.84) |
| | | Eastern & Northern Europe | 0.10 (-0.10, 0.32) | 0.19 (-0.08, 0.51) | 0.24 (0.01, 0.50) |
| | | Eastern Asia | 0.11 (-0.12, 0.34) | 0.22 (-0.10, 0.56) | 0.31 (0.04, 0.61) |
| | | Northern America | -0.00 (-0.18, 0.19) | 0.07 (-0.17, 0.36) | 0.31 (0.11, 0.56) |
| | | Southern & Western Europe | 0.13 (-0.13, 0.36) | 0.23 (-0.09, 0.56) | 0.26 (-0.04, 0.66) |
| | | Southern Asia | -0.03 (-0.23, 0.18) | 0.04 (-0.22, 0.35) | 0.36 (0.15, 0.61) |
| | | Sub Saharan Africa | -0.06 (-0.26, 0.15) | 0.02 (-0.24, 0.30) | 0.39 (0.18, 0.60) |
| | Tan spot | Australia and New Zealand | 0.21 (-0.10, 0.54) | 0.27 (-0.19, 0.77) | 0.23 (-0.06, 0.52) |
| | | Central and West Asia and North Africa | 0.17 (-0.08, 0.44) | 0.26 (-0.13, 0.66) | 0.18 (-0.07, 0.42) |
| | | Eastern & Northern Europe | 0.17 (-0.08, 0.44) | 0.29 (-0.10, 0.69) | 0.13 (-0.15, 0.40) |
| | | Latin America and Caribbean | 0.11 (-0.07, 0.29) | 0.18 (-0.09, 0.45) | 0.17 (-0.00, 0.34) |
| | | Northern America | 0.16 (-0.06, 0.40) | 0.27 (-0.08, 0.66) | 0.12 (-0.11, 0.35) |
| | | Southern & Western Europe | 0.18 (-0.07, 0.44) | 0.32 (-0.06, 0.74) | 0.05 (-0.23, 0.33) |
| | Wheat Blast | Latin America and Caribbean | 0.17 (-0.06, 0.41) | 0.26 (-0.09, 0.62) | 0.13 (-0.10, 0.35) |

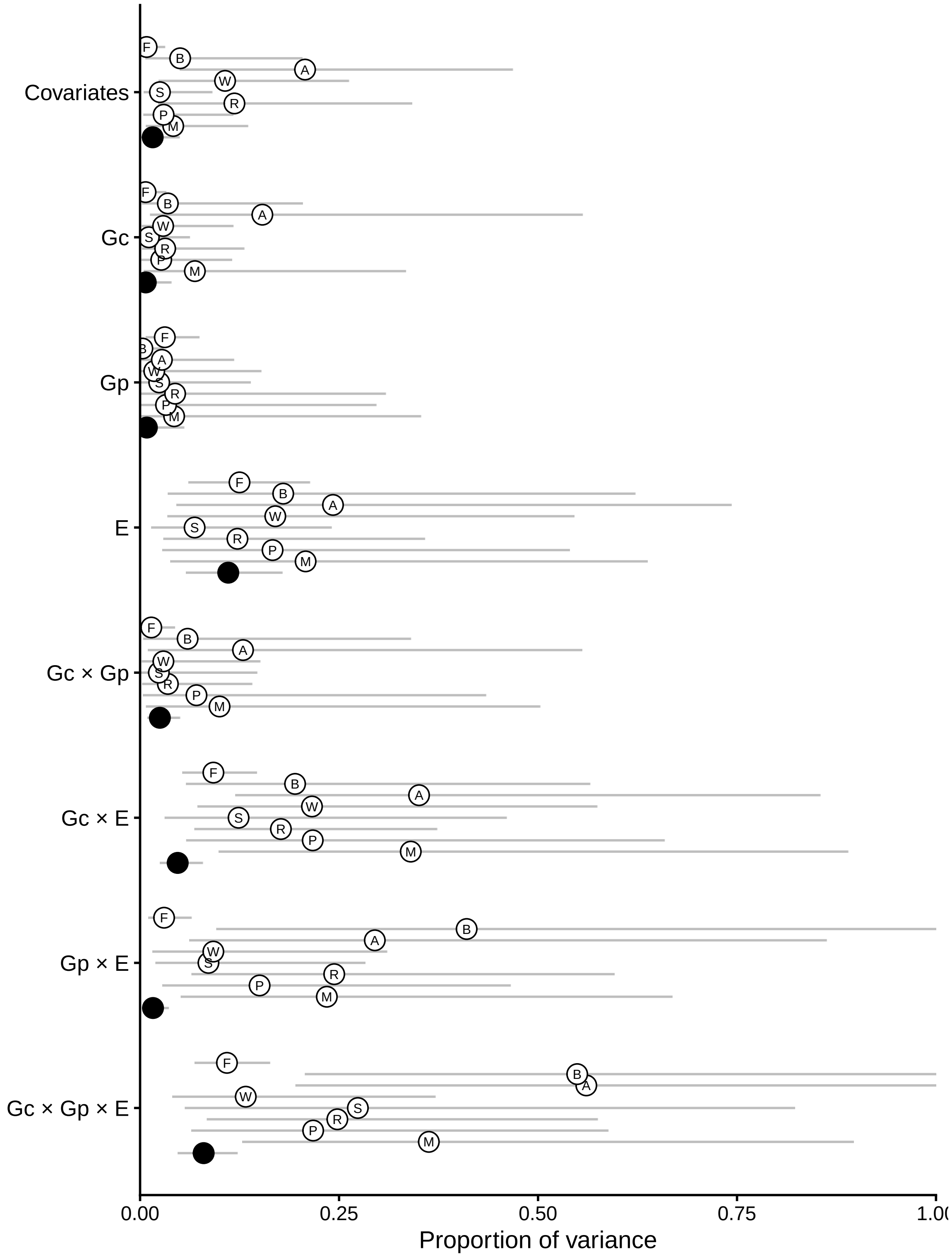


Figure S1: Partitioning of total explained variance in yield loss magnitude by predictor groups for each crop or pest group. This figure is similar to Fig. 1 in the main text, except here each model was fitted to each crop or pest group separately. Key to symbol labels: F = fungi and oomycetes; B = bacteria; A = arthropods; W = wheat; S = soybean; R = rice; P = potato; M = maize.

### Residual diagnosis

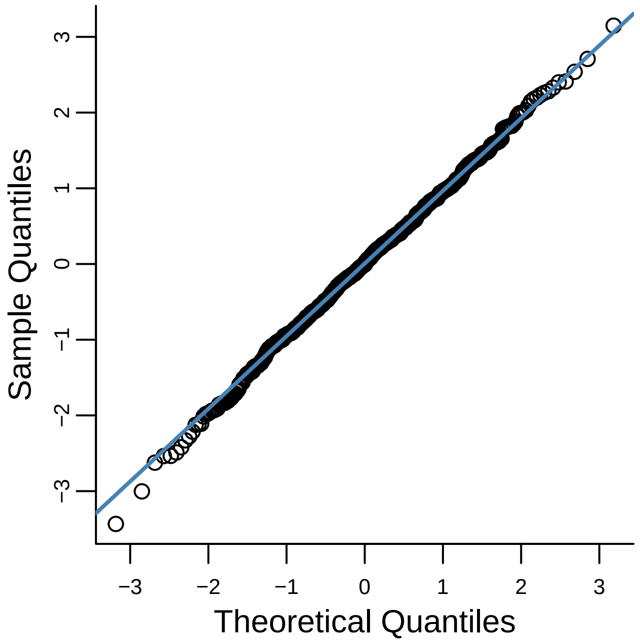


Figure S2: Quantile–quantile plot to diagnose model residuals. The residuals were calculated as randomized quantile residuals following Kay (2023).
